# Quantitatively defining species boundaries with more efficiency and more biological realism

**DOI:** 10.1101/2022.02.14.480439

**Authors:** Jordan Douglas, Remco Bouckaert

**Affiliations:** School of Computer Science, The University of Auckland, Auckland, New Zealand

## Abstract

We introduce a widely applicable species delimitation method based on the multispecies coalescent model that is more efficient and more biologically realistic than existing methods. We extend the phylogenetic tree collapse model to the Yule-skyline model, allowing the ancestral speciation rate to vary through time as a smooth piecewise function. Furthermore, we introduce the cutting-edge proposal kernels of StarBeast3 to this model, thus enabling rapid species delimitation on large molecular datasets and allowing the use of relaxed molecular clock models. We validate these methods with genomic sequence data and SNP data, and show they are more efficient than existing methods at achieving parameter convergence during Bayesian MCMC. Lastly, we apply these methods to two datasets and find inconsistencies with the published literature. Our methods are powerful for rapid quantitative testing of species boundaries in large multilocus datasets and are implemented as an open source BEAST 2 package called SPEEDEMON.

## Main

There are many concepts of what defines a species [1], making species delimitation a field of study that is fraught with pitfalls [2]. Of all the species concepts, the coalescent based species concept is one of the few that allows quantitative testing of different hypotheses [3, 4, 5]. These methods rely on the multispecies coalescent model, where one or more gene trees are constrained within a single species tree [6, 7]. The data used in a multispecies coalescent analysis can consist of multilocus biological sequence alignments, and explicit representations of the gene trees are used in the inference of the species tree, as in the *BEAST [8, 9] model. Alternatively, the data can consist of independently evolving single nucleotide polymorphic (SNP) sites, in which case the gene trees are integrated out [10]. Multispecies coalescent methods can overcome numerous statistical pitfalls underlying traditional phylogenetic analyses which infer species phylogenies from concatenated genomic data [6, 8, 9, 11, 12, 13].

In multispecies coalescent models, the different ways that samples are assigned to species allow us to perform species delimitation in a variety of ways. With Bayes factor delimitation [3, 4] (BFD for gene alignments, BFD* for SNP alignments), hypotheses consist of explicitly stated species assignments. By estimating the marginal likelihood of each of the assignments, the Bayes factor can be estimated in order to compare competing hypotheses in a pairwise fashion. The species tree does not need to be known beforehand, and can be estimated from the data. The methods are implemented in BEAST 2 [14, 15], which means they can be applied with a wide choice of site models, clock models, and tree prior distributions, and combined with a variety of other data, such as morphological features or geographical locations.

An alternative approach is to use reversible jump [16], which allows switching between models during execution of the Markov chain Monte Carlo (MCMC) algorithm where a species is assigned a set of sequences to one where the sequences are split over multiple species, as implemented in BPP [5]. The elegance of this approach is that no explicit sequence assignments to species are required, since these can be either guided through a predefined species tree, or jointly inferred with the species tree. The posterior samples produced by the MCMC algorithm contain a distribution of species assignments from which the various hypotheses under consideration can be tested. Unfortunately, BPP does not support as wide a set of models as BEAST and reversible jump moves are non-trivial to extend for general application to a wide range of models such as optimised relaxed clocks [17].

In contrast, the birth-death collapse model (implemented in DISSECT [18], and STACEY [19]) is a simple but flexible method that does not rely on reversible jump, while still allowing joint inference of sequence assignments to individuals, the phylogeny, and other parameters. First, samples are either given an *a priori* species assignment, or each individual is assigned to its own species. Then, samples whose divergence time falls below some user-defined threshold *ϵ* are considered to be part of the same species, or cluster. This forms the basis of a prior distribution behind the species tree (**Fig. 1**). This spike-and-slab prior is a mixture of a birthdeath tree prior [20] and a collapse model. For nodes above the threshold, only the standard tree prior has an impact (the “slab”), but below the threshold the tree prior is dominated by the “spike”, thus encouraging nodes to remain below the threshold when the user-defined weight of the spike *ω* is large. To this day, the approach is widely applied to species delimitation, and has found its use across a range of taxonomies including amphipods [21], fungi [22], and clingfishes [23].

**Fig. 1:**
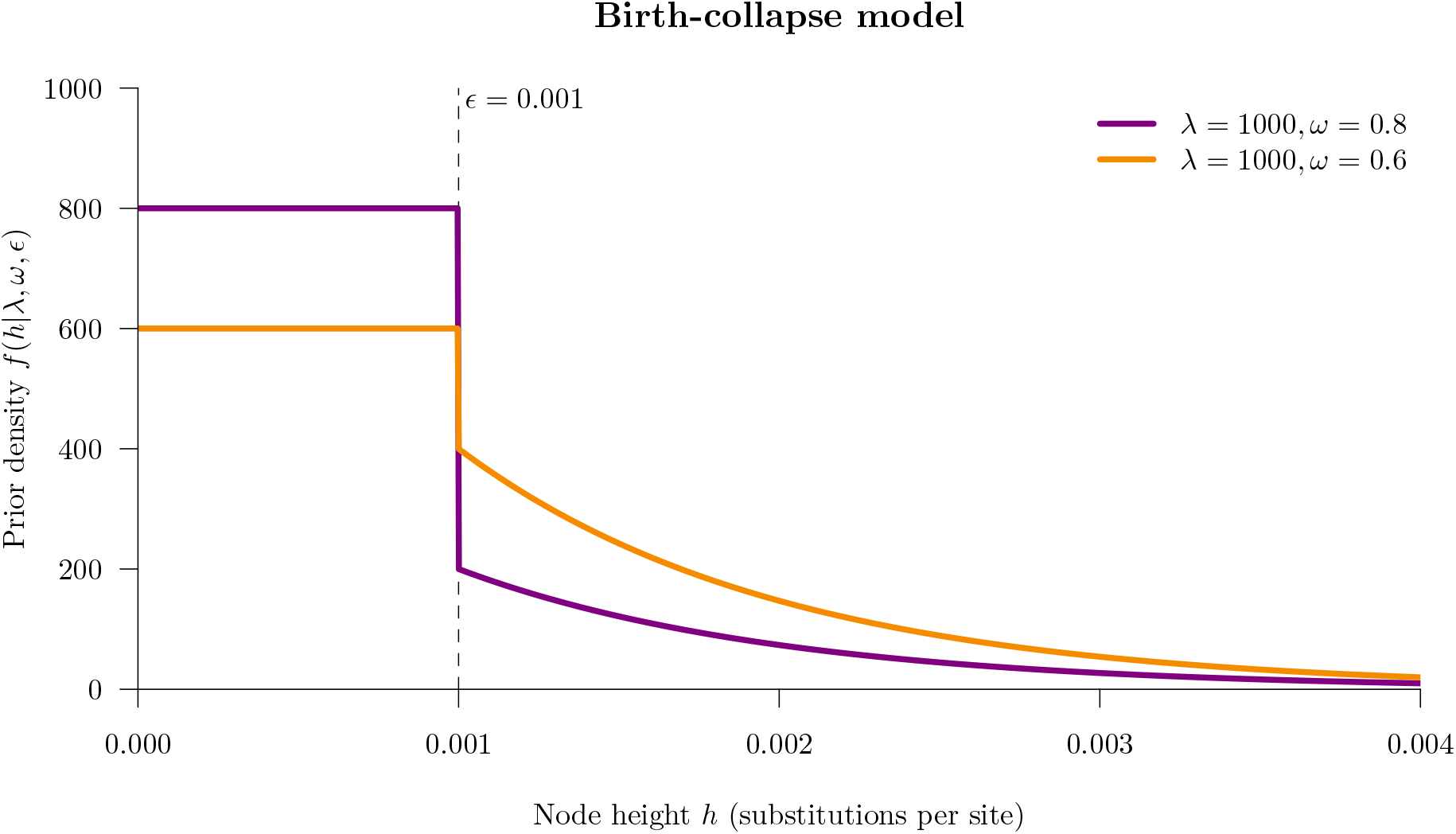
The birth-collapse tree prior distribution with Yule model birth rate *λ* and collapse probability *ω*. Taxa whose common ancestral species lineage falls below threshold *ϵ* are “collapsed” into a single species (or cluster), while species tree nodes above *ϵ* are sampled from an exponential distribution with rate *λ* [24].

Recently, advances have been made in efficient inference under multispecies coalescent models for both gene tree based models (StarBeast3 [25]), and SNP based models (SNAPPER [26]). The threshold approach to species delimitation is readily incorporated into both of these packages as a tree prior distribution.

In this article, we extend the collapse model to allow the speciation rate to vary through time and we demonstrate that this method is a valid approach to performing species delimitation using SNPs with SNAPPER and using gene sequences with StarBeast3. This opens up the way to perform species delimitation in a Bayesian framework using larger datasets and more biologically realistic models compared with previous approaches. We apply these methods to two biological datasets (geckos and primates consisting of lorises and bush babies). Our methods are implemented as the open-source **SPE**ci**E**s **DE**li**M**itati**ON** (SPEEDEMON) package for BEAST 2 [14, 15].

## Results

### Validating the Yule-skyline collapse model

We combined the collapse model [18] with the Yule-skyline model [27] to allow the speciation rate to vary through time as a smooth piecewise function. In this model, the birth rates are analytically integrated and therefore these parameters do not need to be estimated [27]. We call this new tree prior distribution the Yule-skyline collapse (YSC) model.

We validated the YSC model for both SNAPPER (with SNPs) and StarBeast3 (with genes) using a well-calibrated simulation study. In either case, 100 species trees (and their associated gene trees/parameters) were sampled from the prior distribution, and the parameters were recovered using Bayesian MCMC on datasets simulated under the trees. The “true” value of each parameter was compared with the 95% highest density posterior (HPD) interval in order to calculate the coverage. A coverage close to 95% (i.e., from 90 to 99 based on a binomial with p=0.95 and 100 trials) indicates that the model is valid. These experiments suggested that our implementation of the YSC model is valid for the multispecies coalescent. The two well-calibrated simulation studies are presented in **Supplementary Information**.

We also validated these methods for their abilities to identify species assignments, using 100 simulated datasets. To do this, we discretised cluster posterior supports into 20 evenly-spaced bins, and for each bin we counted the number of times each of its clusters existed in the tree from which the data was simulated under. If, for example, a cluster has 52% posterior support, then this hypothesis should be true 50-55% of the time. This experiment confirmed that SNAPPER and StarBeast3 were both able to accurately estimate cluster support probabilities (top panel of **Fig. 2**).

**Fig. 2:**
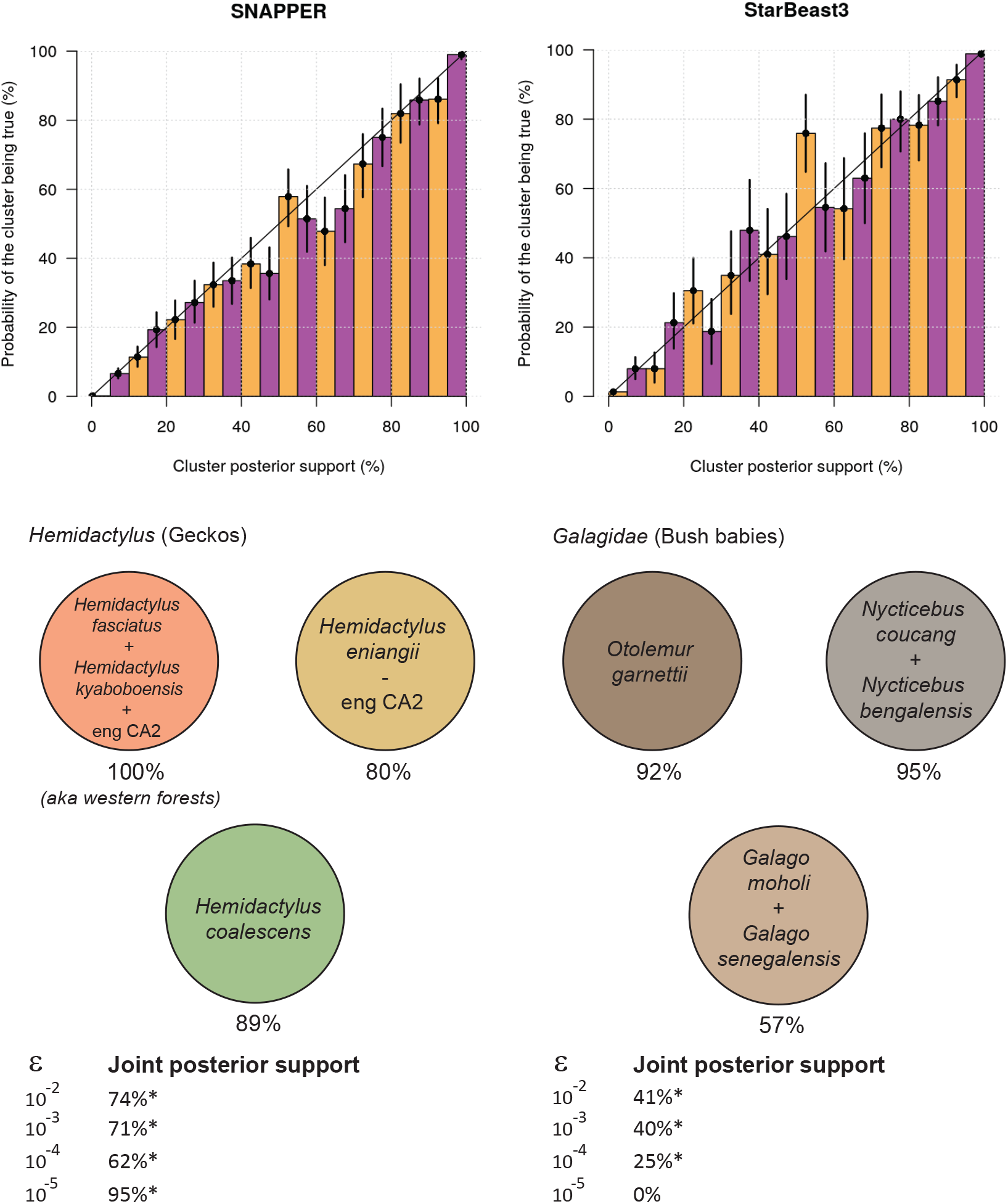
Top: Validating SNAPPER and StarBeast3 for their abilities to recover clusters from simulated data. Each coloured bar has a 95% Binomial credible interval based on the number of clusters used to estimate its probability. Bottom: Species identified in the gecko (left) and primate (right) datasets, under the YSC model. The clustering scheme presented is the one which occurred with the maximal posterior probability across all values of threshold *ϵ* that are labeled with a *. The marginal posterior support is indicated below each cluster (for *ϵ* = 10^-2^). Note that the remaining taxa in the bush baby dataset are omitted from the figure because they exist as singleton clusters.

### Benchmarking the Performance of StarBeast3 and STACEY

We benchmarked the performance of STACEY and StarBeast3 for their abilities to achieve convergence of phylogenetic parameters under the birth-collapse model. To measure convergence, we considered the effective sample size generated per hour of runtime (ESS/hr), for several parameters of interest. Both software packages were benchmarked under the same phylogenetic model, however with effective population sizes analytically integrated by STACEY and estimated by StarBeast3. Although we used a strict clock model here (where the molecular evolution rate is constant through time), we note that StarBeast3 has the further potential of doing inference under the multispecies relaxed clock model [9] using powerful relaxed clock operators [17, 28]. We considered a lizard dataset with 89 samples across 107 loci [29], and a simulated dataset with 48 samples across 100 loci [25]. Each MCMC replicate was run until the effective sample size of the posterior density *p*(*θ|D*) (after a 50% burn-in) exceeded 200.

StarBeast3 outperformed STACEY on both datasets considered (**Fig. 3**). This discrepancy was strongest for the lizard dataset, with StarBeast3 mixing between 1.3 and 9.5 times as fast as STACEY, depending on the parameter, and usually at a statistically significant level. For the simulated dataset, StarBeast3 outperformed in most areas, while STACEY outperformed in others. Most notably, the “slowest” term **min** (i.e., the term which mixed the slowest on any given MCMC replicate) mixed 70% and 120% faster for StarBeast3 on both datasets respectively (*p* < 0.05). Overall, StarBeast3 appears to be the more efficient option.

**Fig. 3:**
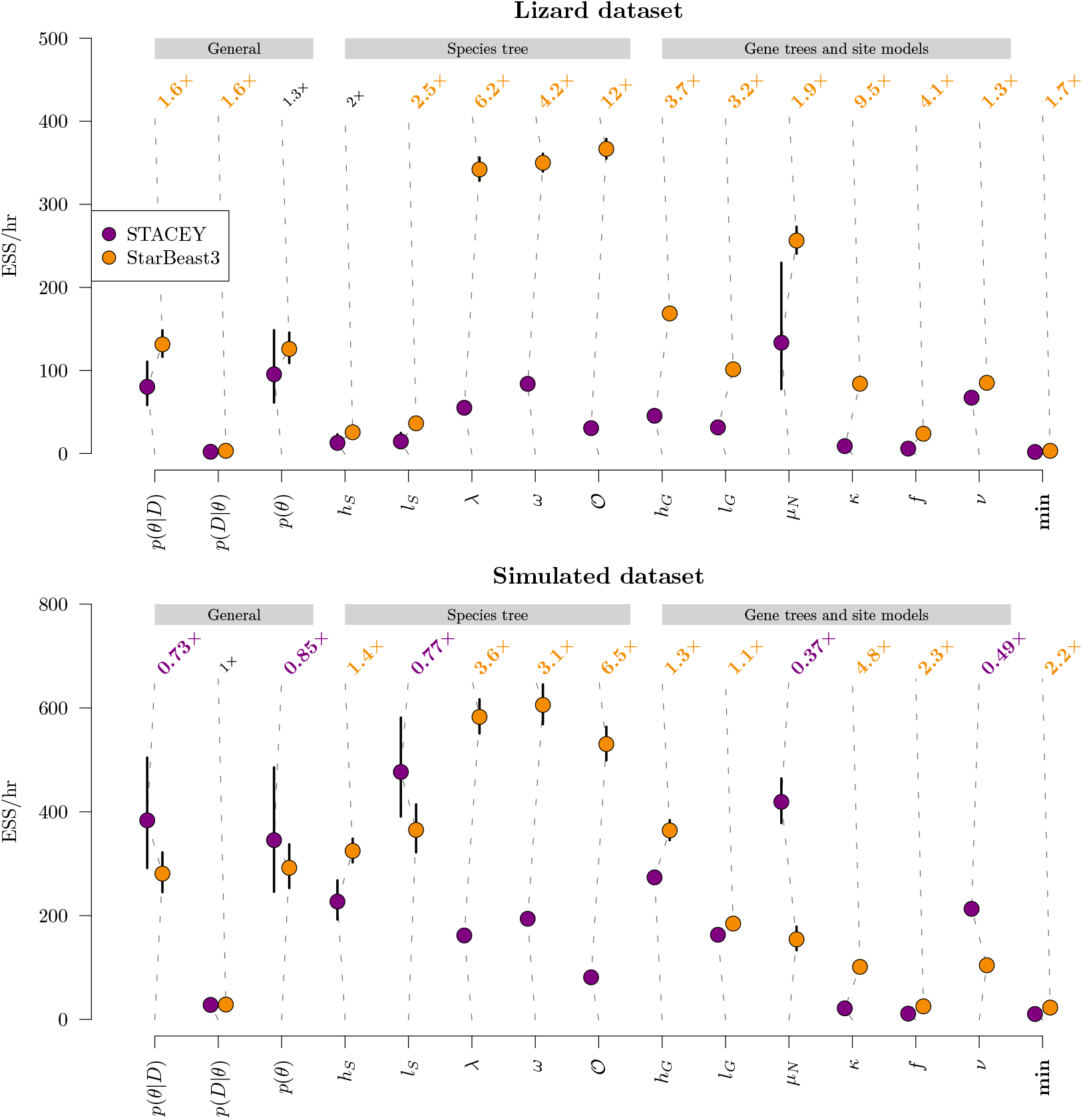
Comparison of parameter exploration efficiencies between STACEY and StarBeast3, under the birth-collapse tree prior. The average effective samples generated per hour (± 1.96 se) is plotted for each term. Top of each plot: a mean relative difference greater (less) than 1 means that StarBeast3 (STACEY) outperformed STACEY (StarBeast3), and is highlighted in orange (purple) font if this difference was significant across 20 MCMC replicates (*p* < 0.05). All means and standard errors were computed in log-space. Notation - *p*(*θ|D*): posterior density; *p*(*D|θ*): likelihood; *p*(*θ*): prior density; *h_S_*: species tree height; *l_S_*: species tree length; *λ*: species tree birth rate; *ω*: species tree collapse weight; 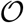: species tree origin; *h_G_*: gene tree height; *l_G_*: gene tree length; *μ_N_*: mean effective population size; *κ*: gene tree transition-transversion ratio; *f*: gene tree nucleotide frequencies; *v*: gene tree substitution rate; **min**: minimum ESS/hr across all other terms.

### Species Delimitation on Gecko SNP Data Using SNAPPER

The *Hemidactylus* are a genus of geckos, found in tropical regions all over the world. To date, there are 180 known species, with newfound species being described every year [30]. Leaché et al. collected 46 samples of genomic data at 1087 loci from 10 forest gecko populations in Western Africa [4, 31]. They identified several species among the populations by explicitly generating multiple species assignment hypotheses (illustrated in Fig. 2 of [4]), and comparing their marginal likelihoods to that of a baseline null hypothesis, using path sampling in conjunction with SNAPP [10] (**Table 1**). This method is known as BFD* and involves one path sampling experiment per hypothesis.

**Table 1:** Comparison of 3 gecko species boundary hypotheses using BFD* and the 129 SNP dataset [4]. For each hypothesis, the number of species, the *log_e_* marginal likelihood ML (averaged across 5 replicates), the Bayes factor BF, and the total rank are reported (with a Yule tree prior). Hypotheses **A** to **F**, were compared by Leaché et al. [4], and **F** ranked the highest. In contrast, **T** was generated by the YSC model presented in this article. These results suggest that **T** is the leading hypothesis, and also demonstrate the power of the collapse model in the task of species delimitation without the need for explicit hypothesis testing.

| Hypothesis | Species | ML | BF | Rank |
| --- | --- | --- | --- | --- |
| A) Baseline | 4 | -1766.1 | — | 3 |
| F) Split *eniangii* | 5 | -1724.3 | 83.6 | 2 |
| T) Lump western with eng CA2 | 3 | -1554.2 | 423.8 | 1 |

Here, we applied the YSC tree prior in conjunction with SNAPPER (instead of SNAPP). In contrast to BFD*, this approach does not require any explicit hypotheses. Instead, we assigned each of the 46 samples to its own species, thus increasing the number of potential hypotheses to *B*_46_ ≈ 2.2 × 10^42^ (Bell number *B*_46_). As a sensitivity analysis, we explored four varying values for threshold *ϵ* = (10^-2^, 10^-3^, 10^-4^, 10^-5^). These results support the lumping of western forest populations into a single species, unlike Leaché et al. (**Fig. 2**). However, these experiments have also identified an individual from the *H. eniangii* population who should have been assigned to the western forest species. Visual inspection of the SNP data also supports this grouping (**Fig. 4**). All four thresholds *ϵ* generated the same leading hypothesis (**Fig. 2**), thus providing high confidence in this species delimitation, and also demonstrating the robustness of this method to varying thresholds.

**Fig. 4:**
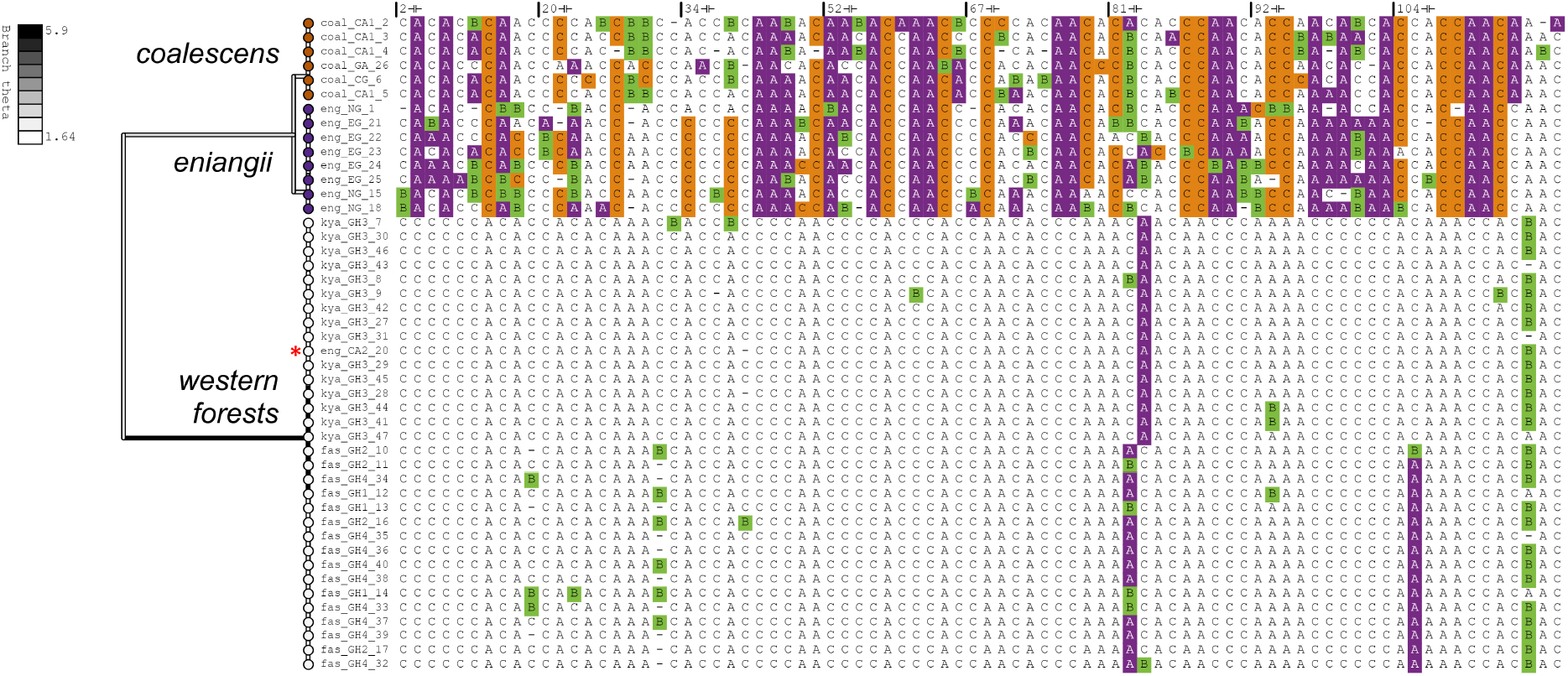
Maximum *a posteriori* species tree of the gecko dataset [4] (left), constructed from 129 genomic loci (right), for *ϵ* = 10^-2^ substitutions per site. Branches are coloured by effective population size *θ*. Only segregating sites (i.e., SNPs which vary among samples) are displayed, and sites are coloured by minor allele (A/C: homozygous; B: heterozygous). The misclassified gecko is indicated by a red asterisk. The analysis was performed using SNAPPER [26] and the figure was generated using PEACH Tree [32].

We denote this newly generated hypothesis as **T**. In order to test **T** (and also to further validate the tree collapse method), we compared it with other hypotheses proposed by Leaché et al., using path sampling (**Table 1**). These results confirmed that **T** is indeed the leading hypothesis, because it had the largest marginal likelihood.

Overall, these experiments have exemplified the major pitfall of the Bayes factor delimitation method: its reliance on explicit species assignment hypotheses. Using this method, we can run a single MCMC analysis and test a large number of hypotheses, whereas BFD* requires a path sampling run for each hypothesis under consideration, and each of these path sampling runs are at least as computationally intensive as a single MCMC run. By using SNAPPER instead of SNAPP, a further order of magnitude in performance gain is accumulated [26].

### Species Delimitation on Bush Baby and Loris Genomic Data Using Star-Beast3

The *Galagidae*, commonly known as the bush babies, and the *Lorisidae* are closely related families of small nocturnal primates [33]. Due to their nocturnal habits, bush babies are fairly understudied compared with other primates and their taxonomy is cryptic [34, 35].

Pozzi et al. compiled a large molecular dataset of the two families and (their outgroups), consisting of 27 genes [34]. We applied the Yule-skyline collapse tree prior, in conjunction with StarBeast3, to infer species boundaries from this dataset. We used the multispecies relaxed clock model to allow substitution rates to vary across lineages [9]. As a sensitivity analysis, we explored four varying thresholds *ϵ* = (10^-2^, 10^-3^, 10^-4^, 10^-5^). Divergence times were calibrated from fossil records, as described by Pozzi et al., and therefore *ϵ* is in units of millions-of-years.

Our resulting phylogeny was in general agreement with that of Pozzi et al. These results contradicted the withstanding taxonomic classifications in three instances (**Fig. 2, 5**). First, two bush baby species (*Galago moholi* and *Galago senegalensis*) were lumped into one (57% posterior support for *ϵ* = 10^-2^). Pozzi et al. hypothesised this contradiction arose as a consequence of taxonomic misclassifications of sequences and/or captive animals. Second, two members of *Galagoides demidoff* were split into two distinct clusters, suggesting that the two individuals might not have belonged to the same species (100% support). This was also reported by Pozzi et al. Finally, two species of the *Lorisidae* were lumped together (*Nycticebus bengalensis* and *Nycticebus coucang*), with 95% support. These three anomalies occurred in the maximal-posterior clustering scheme for three of the four thresholds *ϵ* = (10^-2^, 10^-3^, 10^-4^), thus placing a high level of support in these results, and also demonstrating the robustness of this method to varying *ϵ* (**Fig. 2**). In contrast, *ϵ* = 10^-5^ designated each taxon to its own species (as its maximum *a posteriori* estimate), which is an intuitive result given that *ϵ* = 10^-5^ is equal to just 10 years.

**Fig. 5:**
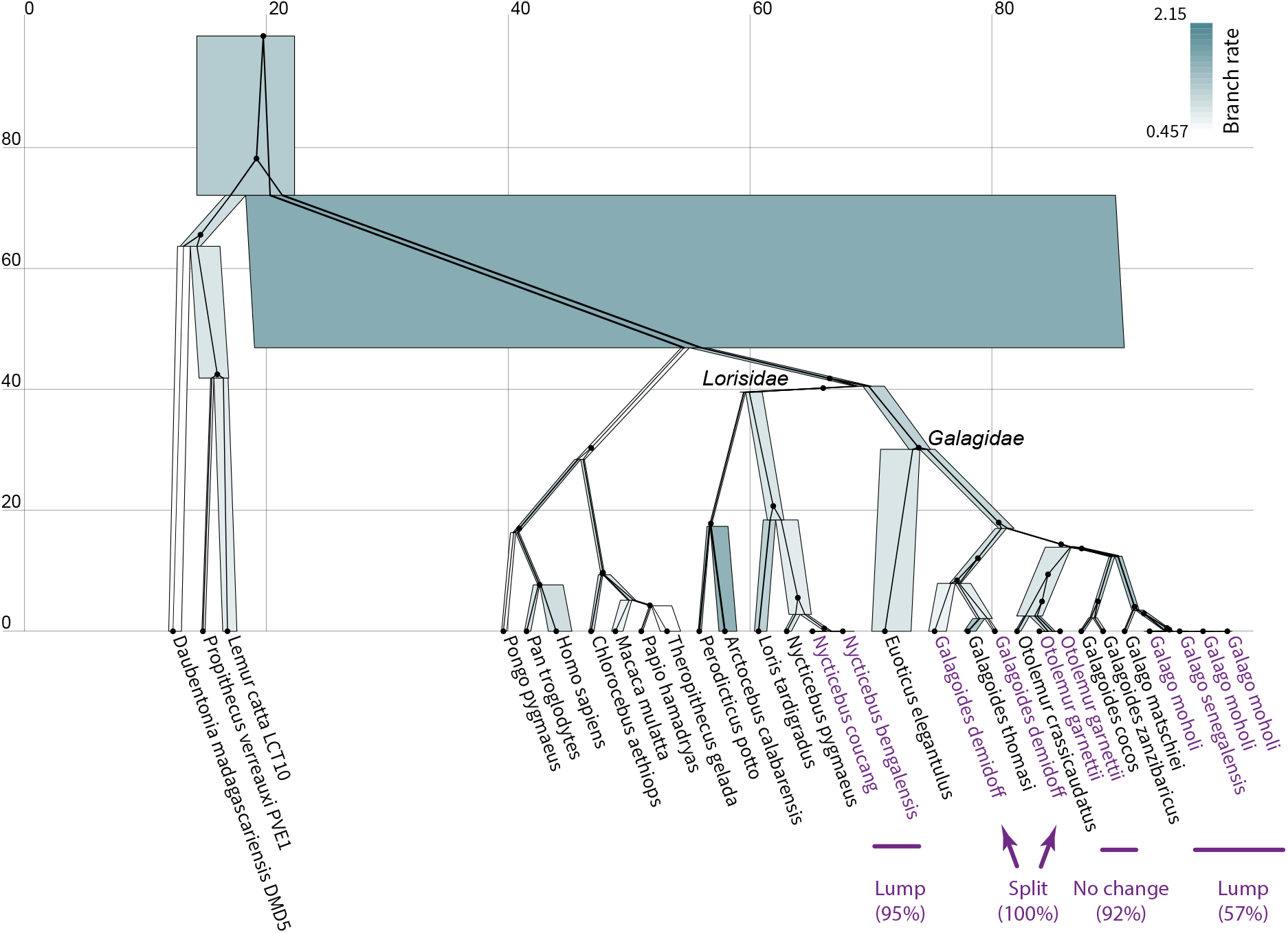
Maximum *a posteriori* species tree of the primate dataset [34]. One arbitrarily selected gene tree (ADORA3) is displayed within the species tree. Node heights are in units of millions of years. Species tree node widths denote effective population sizes, and are coloured by relative branch substitution rates under a relaxed clock model. The posterior support (for *ϵ* = 10^-2^) of the four lump/split/no-change events are displayed. The analysis was performed using StarBeast3 [25] and the figure was generated using UglyTrees [36].

## Discussion

The species delimitation methods we have presented are advanced in both their computational efficiencies as well as their biological realism.

First, we amalgamated the birth collapse model [18] with the Yule-skyline model [27] to enable ancestral speciation rates to vary through time as a smooth piecewise function. In this method, speciation rates are integrated out and the model is reported to converge quite efficiently, despite its increase in complexity over the standard Yule model [24]. Second, we introduced the multispecies relaxed clock model [9] to the species delimitation problem. This model allows molecular evolution rates to vary across species lineages and is therefore more biologically realistic than the withstanding strict clock model. However, these additional complexities in the model are met with highly efficient proposal kernels [28, 17, 25], and much like the Yule-skyline collapse model, is expected to converge quite efficiently in MCMC. Lastly, we demonstrated how the collapse model can be used for molecular sequence analysis in conjunction with StarBeast3 [25] and for SNP analysis in conjunction with SNAPPER [26] - each of which are reported to be significantly more efficient than their predecessors. We demonstrated that StarBeast3 outperforms STACEY at achieving convergence during Bayesian MCMC (**Fig. 3**). We showed how the collapse model can implicitly test all possible species delimitation hypotheses at once (through MCMC), as opposed to one hypothesis at a time (through path sampling; [3, 4]). Overall, these methods are faster and more advanced than other species delimitation approaches.

We validated these advanced methods and applied them to two biological datasets. First, we examined the geckos (genus: *Hemidactylus*) studied by Leaché et al. [4, 31]. Several species delimitation hypotheses were informed by population geography - the leading hypotheses identified 4-5 different species [37]. However, by applying the collapse method to this dataset (without imposing any *a priori* species assignments), we identified an individual from the *H. eniangii* population whose genome was more akin to those from the western forest populations (**Fig. 4**). Our analysis defined 3 species, and the hypothesis was met with high posterior support even across varying collapse model thresholds (**Fig. 2**). It is not immediately clear whether this is a case of taxonomic misclassification, or whether this gecko represents more migration between the forests than anticipated. Although we assigned each sample to its own potential species, it is possible to limit the number of species by for example assigning species to one of six groups such that each of the seven hypotheses considered in the BFD* analysis can be formed by collapsing the species tree. However, this would not have allowed us to find the best fitting assignment, because the misclassified sequence eng CA2 20 would not be allowed to cluster with the western forest sequences. Therefore, we recommend assigning each sample to its own species when computationally feasible.

Second, we examined the primates (families: *Galagidae* and *Lorisidae*) studied by Pozzi et al. [34]. We showed that four bush babies should have been lumped into a single species, instead of two (*Galago moholi* and *Galago senegalensis*), and we identified a paraphyletic relationship between two members of *Galagoides demidoff*. Both observations have a moderate-to-high level of posterior support, across a range of collapse thresholds (**Fig. 2**), and we therefore concur with Pozzi et al. Our analysis also lumped two further *Lorisidae* species together (*Nycticebus bengalensis* and *Nycticebus coucang*) with 95% posterior support, thus providing high confidence that these two individuals were in fact from the same species.

For both datasets considered, the collapse model unveiled anomalies underpinning their taxonomic classifications. It is indeterminate from genomic data alone whether these are trivial labeling errors (at the sequence level or at the animal level) or whether they represent nontrivial biological processes. Either way, automated methods like this one, that make no *a priori* assumptions about species assignments, can remove some of the burden from the researcher carrying out such analyses.

The methods discussed here can be further advanced by reducing the size of the search space. When ancestral relations among a set of taxa are firmly established, a fixed topology analysis may be sufficient. In this case, the species tree topologies can be fixed at some non-disputed estimate, with only their node heights, and therefore species boundaries, estimated during MCMC. This would reduce the search space and further expedite the analysis. Alternatively, the species boundary hypothesis space can be restricted without the need to fix the topology or generate explicit hypotheses. This can be achieved by introducing monophyletic constraints onto the species tree. Both of these scenarios are readily achieved in BEAST 2 and the collapse tree prior is applicable in either case.

However, the methods discussed in this article come with their limitations. First, the collapse model is reliant on a threshold parameter *ϵ*, and it is not clear what this threshold should be. Although there is a moderate degree of robustness to this term (**Fig. 2**, and [18]), it would be beneficial to have a method which explicitly estimates the species assignment function without the need for such a heuristic. However, such an improvement may be met with convergence difficulties during Bayesian MCMC. Second, the collapse model is not applicable to ancestral lineages. Lineages which date back before the threshold *ϵ* (including ancestral samples) are unable to be clustered under the collapse model in its current form. Further, as pointed out by Jones et al. [18], the multispecies coalescent model has assumptions such as lack of hybridisation that are likely to be violated and may impact the results of the species delimitation analysis. The method does not correct cluster bias due to sampling selection bias and its behaviour with ring species is unclear.

## Conclusion

The collapse model is a phylogenetic tree prior distribution (**Fig. 1**) used for species delimitation under the multispecies coalescent [18]. We advanced the work by Jones et al. by formally validating this method through well-calibrated simulation studies (**Fig. 2** and **Supplementary Information**), and we demonstrated that the recently developed StarBeast3 [25] and SNAPPER [26] inference engines outperformed existing methods at the task of fast Bayesian species delimitation (**Fig. 3**). Furthermore, we combined the collapse model with our Yule-skyline model [27] to allow the species tree birth rate to vary as a smooth piecewise function over time. We applied the Yule-skyline collapse model to two biological datasets; gecko SNP data [4] and primate genomic data [34]. In either case, we identified species boundaries that contradicted those assigned to individuals in the original datasets (**Fig. 4 – 5**), thus exemplifying the appeal of the method.

The methods presented are implemented in the SPEEDEMON package for BEAST 2 and are suitable for rapidly identifying species on large datasets with over 100 genes or thousands of SNPs. The implementation in BEAST 2 allows adding various other types of data to the species tree, such as morphological features (as recommended by Olave et al. [38]) and geographical location [39, 40]. Together, SPEEDEMON provides a flexible package for species delimitation catering to a wide range of biological applications.

## Methods

### Collapse models

Let *T* be a binary rooted time tree over *n* taxa with leaf nodes *x*_1_,…, *x_n_* and internal nodes *x*_*n*+1_,…, *x*_2*n*-1_. Let *h_i_* ≥ 0 denote the height of node *i*, where all leaves are assumed to be extant with height *h_i_* = 0. Suppose, we have a distribution over trees *f*(*T*|*θ*) for some set of parameters *θ*, such as a Yule or birth death distribution, where *f* can be written as the product of internal node height contributions. That is, we can write *f*(*T|θ*) as 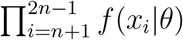. Furthermore, we assume that *f*(*x_i_|θ*) = 1 if *h_i_* = 0, so internal nodes of height zero do not contribute to this tree distribution. To avoid numerical instabilities associated with zero-node-heights, we will assume that nodes below some threshold *ϵ* do not contribute to the branching/coalescent process, and *f*(*T|θ, ϵ*) = ∏_*n*≤*i*≤2*n*-1,*h_i_*≥*ϵ*_ *f*(*x_i_*|*T, θ*) where *f*(*x_i_|T,θ*) = 0 for *h_i_* < *ϵ*.

Now, let us define the collapse tree prior as the weighted sum of some tree distribution *f*(*T|θ, ϵ*) with a spike density *m*(*x_i_|ϵ*) on internal nodes heights, where *m*(*x_i_|ϵ*) = 0 if *h_i_* > *ϵ* and *m*(*x_i_|ϵ*) = 1/*ϵ* otherwise (**Fig. 1**). Let *ω* be a weight between 0 and 1 that governs the contribution of the components of the mixture. Then, the collapse tree prior *f*(*T|θ, ϵ, ω*) can be written as

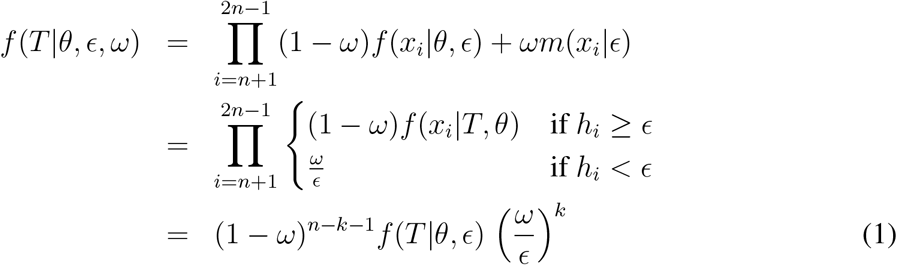

where *k* is the number of internal nodes with node heights less than *ϵ*. In this study, we fixed *ϵ* to a small, e.g., 10^-4^ substitutions per site, and sampled the value of *ω* during MCMC.

When using the birth-death distribution as a tree distribution *f*(*T|θ,ϵ*), we get the birthdeath collapse model defined for DISSECT and STACEY [18, 19]. This model is conditioned on an origin height and its parameters *θ* consist of a birth rate, a death rate, and the origin height. By setting the death rate to zero, the widely-used Yule model is obtained [24].

Alternatively, we can use the Yule skyline model [27], which is a pure birth model that conditions on the number of extant species *n* – *k* – 1. This model splits up time into epochs and can therefore be naturally extended to the case where nodes are collapsed below *ϵ* height. The Yule skyline model integrates out the birth rate skyline (which is assumed to follow a gamma prior), and allows the smoothing of birth rates over epochs, where the birth rate prior at epoch *i* + 1 is conditional on the birth rate posterior estimate at epoch *i*. In this model, *θ* consists of the shape and rate of the gamma prior of the first epoch. This forms the basis for the Yule-skyline collapse (YSC) model.

Another suitable epoch model is the birth-death skyline model [41], which allows different birth rates and death rates in each epoch, and can easily be adapted to ignore events in the epoch with height less than *ϵ*. While the Yule model assumes all species are observed, the birth-death skyline model introduces a sampling proportion parameter *ρ*. In general, any tree distribution that can be decomposed into contributions of the individual nodes in the tree can be combined with the collapse model, for instance, the multi type tree distribution [42] allows rate changes at arbitrary locations in the tree.

### Prior Distributions

For SNAPPER, we used the YSC tree prior with coalescent rates ~ Gamma(*α* = 100, *β* = 0.01) and collapse weight *ω* ~ Beta(*α* = 1, *β* = 2) under the prior distribution. The skyline consists of 4 epochs, where the birth rate of the first epoch is drawn from a Gamma(*α* = 2, *β* = *b*) prior where *b* ~ Log-normal(*μ* = 1.59, *σ* = 0.2).

For STACEY, we used the strict clock model and the birth-collapse tree prior with collapse weight *ω* ~ Beta(*α* = 1, *β* = 1), birth rate *λ* ~ Log-normal(*μ* = –2.43, *σ* = 0.5), and origin height 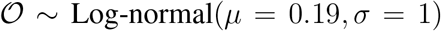 under the prior distribution. Species tree branchwise effective population sizes were drawn from an Inverse-gamma distribution with a shape of 2, and a mean of *μ_N_*, where *μ_N_* ~ Log-normal(*μ* = 2.87, *σ* = 0.5). Nucleotide evolution was assumed to follow an HKY substitution model [43] with transition-transversion ratio *κ* ~ Log-normal(*μ* = 1, *σ* = 1.25), nucleotide frequencies *f* ~ Dirichlet(*α* = (10,10,10,10)), and substitution rate *v* ~ Log-normal(*μ* = –0.18, *σ* = 0.6). Each gene tree was associated with an independent and identically distributed substitution model.

For StarBeast3, we used the same model as STACEY during performance benchmarking (but with effective population sizes estimated instead of integrated out). However for the general case, we instead ran StarBeast3 under the YSC species tree prior (with 4 epochs) and the multispecies relaxed clock model [9], with species branch rates drawn from 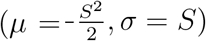 and standard deviation *S* ~ Gamma(*α* = 5, *β* = 0.05).

### Proposal Kernels

We employed the proposal kernels of SNAPPER, STACEY, and StarBeast3 when doing inference under the collapse model. When benchmarking StarBeast3 against STACEY, StarBeast3 gene tree inference was parallelised with 4 threads while STACEY was run with just 1 thread (as it does not possess any equivalent benefit from multithreading).

We also introduce one further tree node height operator which increases or decreases the number of clusters in the species tree. This operator is known as `ThresholdUniform` and works as follows:

1. Sample *B* ~ Bernoulli(0.5).
2. If *B* = 0, then let *x* be an internal node from *T* such that *h_x_* ≥ *ϵ*, and *h_l_, h_r_* < *ϵ*, where *l* and *r* are the children of *x*. Let the lower limit *t*_0_ = max{*h_l_, h_r_*} and let the upper limit *t*_1_ = *ϵ*.
3. If *B* = 1, then let *x* be an internal node from *T* such that *h_x_* < *ϵ*, and *h_p_* ≥ *ϵ*, where *p* is the parent of *x*. Let the lower limit *t*_0_ = *ϵ* and let the upper limit *t*_1_ = *t_p_*.
4. If there are no such eligible nodes for *x*, then reject the proposal.
5. Propose a new value for *h_x_* as: 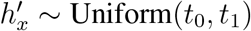.

This proposal adjusts the height of a species tree internal node from one side of the threshold boundary (at height *ϵ*) to the other. This operation will either lump two clusters together or split one cluster into two, without affecting the species tree topology.

## Software Availability

SPEEDEMON is available as an open-source BEAST 2 package with an easy-to-use graphical user interface. Instructions for downloading and running SPEEDEMON can be found at https://github.com/rbouckaert/speedemon.

## Supplementary Data

Well-calibrated simulation study results can be found in SI Appendix. BEAST 2 XML files used in this study can be found at https://github.com/jordandouglas/speedemon_SI.

## Acknowledgements

The study was supported by a Marsden grant 18-UOA-096 from the Royal Society of New Zealand. Software packages were benchmarked using the New Zealand eScience Infrastructure (NeSI) cluster, funded by the New Zealand Ministry of Business, Innovation and Employment.

## Notes

https://github.com/rbouckaert/speedemon

### Competing Interest Statement

The authors have declared no competing interest.

