## Appendix 1 for "Quantitatively defining species boundaries with more efficiency and more biological realism"

February 15, 2022

### Supplementary Information

Our two well-calibrated simulation studies are presented in the figures below. The coverage of each parameter is close to 95% thus providing confidence in the validity of these methods.

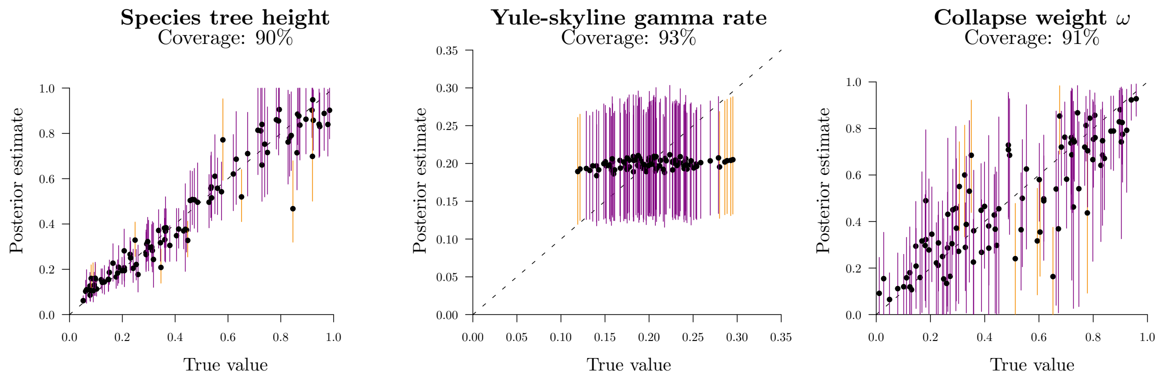


Fig. S 1: Well-calibrated simulation study of the YSC model under a multispecies coalescent framework, with gene trees integrated out (SNAPPER). The species tree consisted of 40 taxa, and four samples were assigned to each species tree taxon *a priori*. Each of the 100 gene trees are associated with one SNP. The mean parameter estimates (under the posterior distribution) are indicated by black circles, and 95% highest posterior density (HPD) intervals are coloured dark if the true value is in the interval, or light otherwise. Most of the terms have close to 95% coverage under the 95% HPD, therefore suggesting that the model is both valid and correctly implemented. Omitted from this figure are the coalescent rates; which also have close to 95% coverage. “True” values were sampled from the joint prior distribution using MCMC.

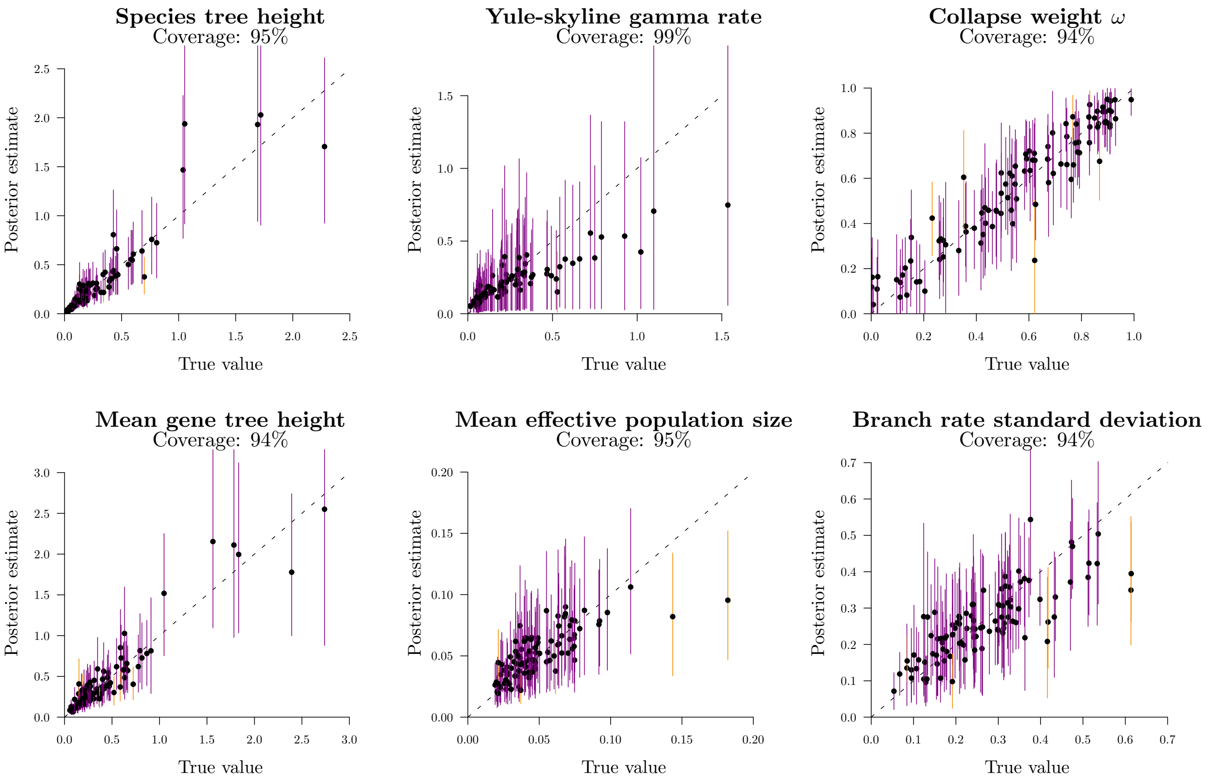


Fig. S 2: Well-calibrated simulation study of the YSC model under a multispecies coalescent framework, with gene trees estimated (StarBeast3). The species tree consisted of 40 taxa, and one sample was assigned to each *a priori*. Each gene tree was inferred from a respective nucleotide sequence 0.5kb in length. Mean gene tree heights were averaged across 4 gene trees, and the multispecies relaxed clock model was applied (Ogilvie et al., 2017). These results suggest that the model is likely to be both valid and correctly implemented (see Fig. S 1 for figure notation). Omitted from this figure are the gene-tree HKY substitution model parameters; which also have close to 95% coverage. Parameters were directly sampled from the prior in order to capture the generative process.
